# Observing the suppression of individual aversive memories from conscious awareness

**DOI:** 10.1101/2021.10.17.464746

**Authors:** Xuanyi Lin, Danni Chen, Ziqing Yao, Michael C. Anderson, Xiaoqing Hu

## Abstract

When reminded of an unpleasant experience, people often try to exclude the unwanted memory from awareness, a process known as retrieval suppression. Despite the importance of this form of mental control to mental health, the ability to track, in real time, individual memories as they are suppressed remains elusive. Here we used multivariate decoding on EEG data to track how suppression unfolds in time and to reveal its impact on cortical patterns related to individual memories. We presented reminders to aversive scenes and asked people to either suppress or to retrieve the scene. During suppression, mid-frontal theta power within the first 500 ms distinguished suppression from passive viewing of the reminder, indicating that suppression rapidly recruited control. During retrieval, we could discern EEG cortical patterns relating to individual memories-initially, based on theta-driven, visual perception of the reminders (0-500 ms) and later, based on alpha-driven, reinstatement of the aversive scene (500-3000 ms). Critically, suppressing retrieval weakened (during 420-600 ms) and eventually abolished item-specific cortical patterns, a robust effect that persisted until the reminder disappeared (1200-3000 ms). Actively suppressing item-specific cortical patterns, both during an early (300-680 ms) window and during sustained control, predicted later episodic forgetting. Thus, both rapid and sustained control contribute to abolishing cortical patterns of individual memories, limiting awareness, and precipitating later forgetting. These findings reveal how suppression of individual memories from awareness unfolds in time, presenting a precise chronometry of this process.

## Introduction

Following an upsetting event, memories of the experience often come to mind uninvitedly. Even seemingly innocuous reminders can bring us back to the traumatic scene in the blink of an eye, triggering intrusive memories and distress. When this happens, people often recruit inhibitory control to terminate unwelcome retrieval, a process known as retrieval suppression (Anderson & Hulbert, 2020; Küpper et al., 2014). An ability to control aversive memories and to keep them out of awareness can promote resilience and safeguard mental well-being, especially in the aftermath of trauma (Anderson & Hanslmayr, 2014; Catarino et al., 2015; Engen & Anderson, 2018; Hu et al., 2017; Mary et al., 2020). Despite the fundamental importance of this process, much remains unknown about its basic mechanisms. Indeed, no study has yet observed individual memories as they are suppressed, a pre-requisite to tracking the dynamics of memory control. Observing suppression unfold in real time is fundamental to advance neurobiological models of memory control, and to inform novel interventions that may aid people in forgetting unwanted memories.

Neuroimaging research suggests that during retrieval suppression, when a person sees a reminder to an unwanted memory, the prefrontal cortex exerts inhibitory control over the hippocampus and its adjacent medial temporal lobe structures to stop retrieval (Anderson et al., 2004; Depue et al., 2007). Furthermore, inhibitory control down-regulates activity in content-specific neocortical areas implicated in the encoding of the original memory (Benoit et al., 2015; Depue et al., 2007; Gagnepain et al., 2014, 2017; Hu et al., 2017; Mary et al., 2020). Given its limited temporal resolution, however, functional magnetic resonance imaging does not permit a detailed account of the temporal dynamics underlying the suppression of individual memory representations.

Conversely, although EEGs have the temporal resolution needed to track the online dynamics of retrieval suppression (Bergström et al., 2009; Hellerstedt et al., 2016; Hu et al., 2015; Zhang et al., 2016), its poor spatial resolution has historically rendered it difficult to isolate individual memories as they are suppressed. However, advances in multivariate pattern analysis have allowed researchers to exploit distinctive EEG scalp distributions to identify specific memory representations (Bae & Luck, 2018; Treder et al., 2021; Wolff et al., 2017). Here, leveraging EEGs’ temporal resolution, and multivariate decoding analyses, we sought to isolate cortical EEG patterns unique to individual memories, and to observe suppression abolishing such patterns in real time. For this purpose, we adapted the think/no-think paradigm to require our participants to voluntarily retrieve or to suppress aversive scenes when confronting reminders (Anderson & Green, 2001; Depue et al., 2007; Küpper et al., 2014). To track the temporal dynamics of retrieval suppression, we took a twostep approach to our EEG analysis. First, we used decoding to determine how and when suppression differed, in general, from retrieval; thus, using data from all EEG sensors, we applied multivariate EEG analysis to compare retrieval and retrieval-suppression manipulations to a perceptual baseline condition, in which neither retrieval nor suppression had occurred. Pairwise condition-level decoding should reveal neural dynamics of retrieval and retrieval suppression, relative to the no-retrieval baseline. We focused on the role of frontal theta within the first 500 ms, given frontal theta power increase has been related to inhibitory control processes (Anderson & Hulbert, 2020; Cavanagh & Frank, 2014; Crespo-García et al., 2021; Nigbur et al., 2011).

We next used MVPA within each condition to isolate item-specific cortical EEG patterns and to examine their development over time in relation to the suppression process. We hypothesized successful suppression and forgetting of unwanted memories involves two key requirements. First, inhibitory control needs to act rapidly to truncate retrieval before the reminder elicits episodic recollection, reinstating the aversive scene. Second, inhibition must be sustained over time and expunge intruding memories from awareness, abolishing residual cortical reinstatements. The initial truncation of retrieval must proceed very rapidly; research on the time course of memory retrieval reveals a chronometry with multiple stages. Upon visually perceiving a memory cue, a cue-to-memory conversion process is thought to occur within 500 ms, along the occipital-temporal cortex pathway. Outputs of this process are thought to arrive in the hippocampus, initiating pattern completion at around 500 ms. Pattern completion is thought to then drive cortical reinstatement of the associated target memory during the 500-1500 ms window, at least for simple laboratory materials (Staresina et al., 2019; Staresina & Wimber, 2019; Treder et al., 2021; Yaffe et al., 2014).

Based on these findings, we hypothesized that countering the emergence of item-specific cortical patterns would involve inhibitory control to target processes in the cue-to-memory conversion window (at around 500 ms) and also in the cortical reinstatement window (500-1500 ms). To understand how suppression modulates item-specific activity, we further examined 4-8 Hz theta activity during the early 0-500 ms time window, given the roles of theta in sensory intake and feedforward information flow originating from the sensory cortex (Bastos et al., 2015; Colgin, 2013). To track reinstatement, we focused on 9-12 Hz alpha activity in the 500-1500 ms window, given alpha activity’s role in working memory maintenance and reinstatement (Fellner et al., 2020; Jensen et al., 2002). By comparing how item-specific cortical patterns unfold over time between the retrieval and the retrieval suppression conditions, we gain a window into the timeline for how inhibitory control affects the recollection of individual memories.

We found that suppressing retrieval enhanced early theta control and began to attenuate item-specific cortical patterns within the first 500 ms, likely disrupting the perception-to-memory conversion processes. Critically, retrieval suppression weakened and ultimately abolished item-specific cortical patterns during the 500-1500 ms memory reinstatement window in a sustained manner. These results were especially pronounced among participants who successfully forgot the unpleasant scenes that they suppressed; in contrast, less successful forgetting was associated with insufficient mobilization of early theta control mechanisms, and relapse of cortical patterns for unwelcome content during the full suppression window.

## Results

### Suppressing Retrieval Induces Forgetting of Emotional Memories

Following the emotional Think/No-Think (TNT) task, participants completed a cued recall test during which they verbally described the aversive scene that they thought was linked to each of the cue objects. We coded and scored verbal descriptions on *Identification, Gist* and *Detail* (see Methods). Each of these three scores was submitted to a one-way repeated-measure (Think, No-Think and Baseline) analysis of variance (ANOVA). Results showed a significant condition effect on *Identification F* (1.87,72.93) = 7.35, *p* = .002; *Detail* (*F* (1.93,75.2) = 13.79, *p* < .001 and *Gist* (*F* (1.92,74.95) = 6.22, *p* = .004). Planned contrasts comparing Baseline and No-Think conditions confirmed that participants showed significant suppression-induced forgetting on *Identification, t*(39) = -2.07, *p* = .045, *dz* = 0.33, and *Details, t*(39) = -2.16, *p* = .037, *dz* = 0.34, whereas the forgetting effect on *Gist* was not significant *t*(39) = -1.58, *p* = .123, *dz* = 0.25, see Figure 1B).

**Fig. 1.**
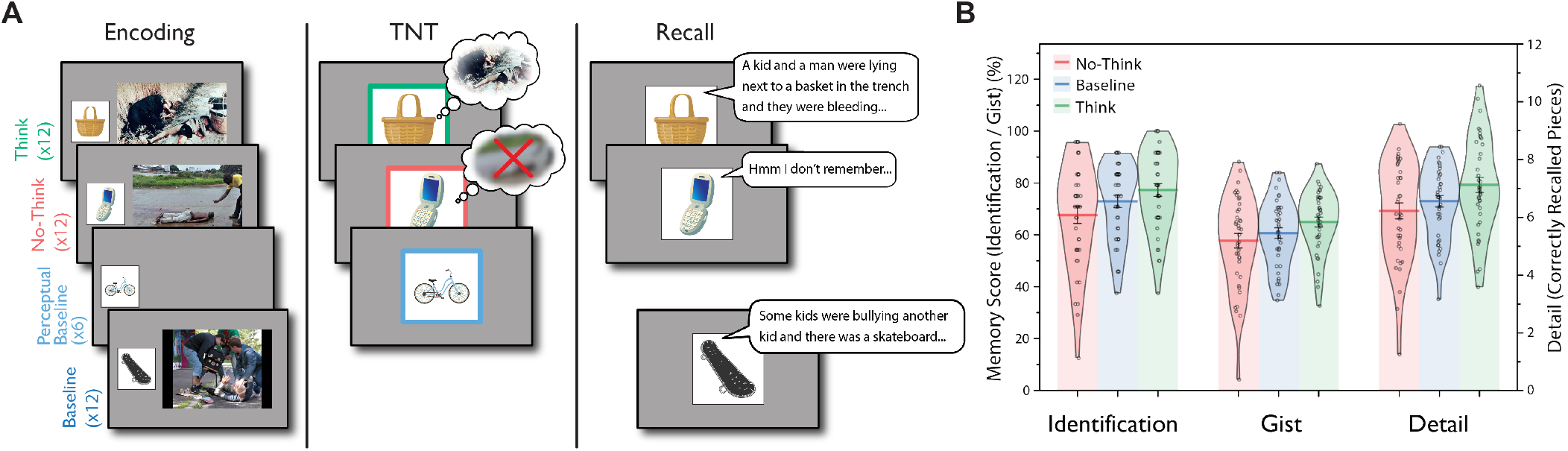
Experimental Procedure and Suppression-Induced Forgetting. (A) The emotional Think/No-Think task (eTNT) included three phases. 1) Encoding: Participants first learned object-aversive scene stimulus pairings; and they also viewed some objects without any paired scene (i.e., Perceptual Baseline); 2) Think/No-Think (TNT) task: Participants either retrieved (Think) or suppressed the retrieval (No-Think) of negative scene memories. Participants were also presented with Perceptual Baseline trials without any retrieval; Think, No-Think, and Perceptual Baseline instructions were cued by a green, red, or blue colored box respectively, surrounding the cue object; 3) Recall: Participants viewed object cues and verbally described their associated scenes. (B) Suppression-Induced Forgetting on Identification, Gist and Detail measures from the Recall test. Suppression-induced forgetting can be seen in the lower recall of No-Think than Baseline items (n = 40).

### Stopping Retrieval is Distinct From Not-Retrieving

We next sought to identify EEG activity tied to stopping retrieval. Towards that end, we examined EEG activities that distinguished No-Think, Think, and Perceptual Baseline (i.e., no-retrieval) conditions. In the time domain, condition-level multivariate decoding not only distinguished retrieval suppression from voluntary retrieval (NT vs. T, *p*_corrected_ < .001, Figure 2C, purple), but also from non-retrieval in our perceptual baseline condition (NT vs. PB, *p*_corrected_ < .001, Figure 2C, red). These differences imply that unique cognitive operations contributed during retrieval suppression, consistent with the involvement of an active stopping mechanism. Differences between NoThink and Think conditions emerged as early as 140 ms and persisted throughout the entire trial period until ∼3000 ms. In addition, we also could distinguish retrieval from non-retrieval (T vs. PB, *p*_corrected_ < .001, Figure 2C, green). At least some of the latter decoding difference arose from EEG correlates of active retrieval processes during the Think condition: decoding accuracies from 500-3000ms during the Think vs. Perceptual Baseline analysis predicted retrieval-induced facilitation of Think items in the Identification measure, r = 0.33, *p* =.036; and in the Detail measure, r = 0.33, *p* =.041 (Figure 2D, also see Figure S2A).

**Fig. 2.**
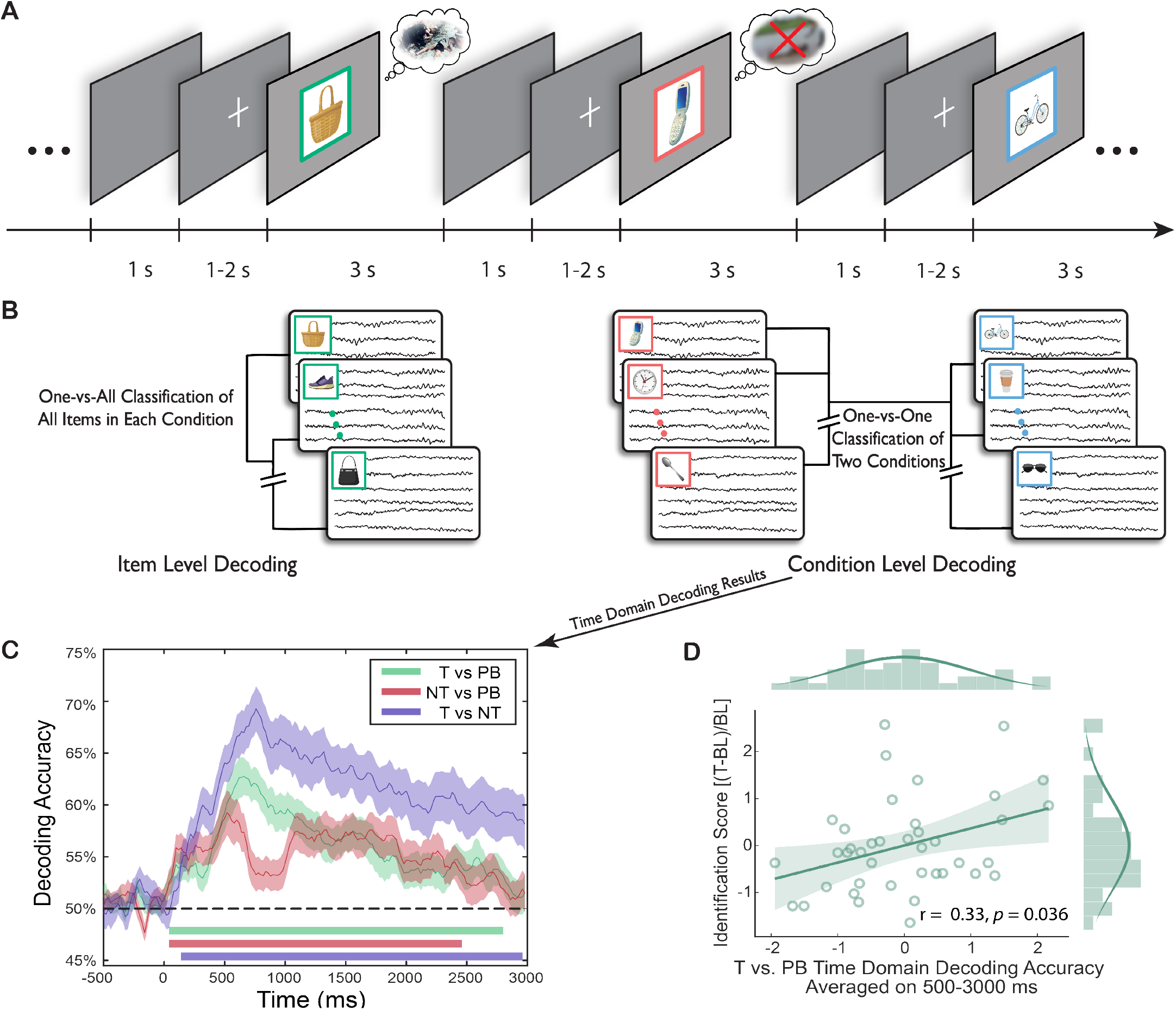
Decoding Approaches Diagram and Condition-level Time-domain EEG Decoding Results. (A-B) An illustration of trial flow in the EEG-based eTNT task, and the logic of decoding analyses. (C) Condition-level decoding based on time domain EEGs revealed significant differences in all three pairwise comparisons. Colored lines along x-axis indicate significant clusters (permutation cluster corrected): No-Think vs Perceptual Baseline, 40-2460 ms, *p*_corrected_ < .001; Think vs Perceptual Baseline, 40-2800 ms, *p*_corrected_ < .001; Think vs No-Think, 140-2960 ms, *p*_corrected_ < .001. Shaded areas indicate standard errors of the mean (S.E.M). (D) Time domain Think vs. Perceptual Baseline decoding accuracies during the 500-3000 ms window was positively correlated with the enhancement of Think item recall on the final recall test, on the Identification score ((Think –Baseline)/Baseline, or the recall benefit, proportional to baseline).

Retrieval suppression could also be distinguished from retrieval and passive viewing based on time-frequency domain EEGs. Between condition decoding revealed differences among all pairwise comparisons (Figure 3A-F). Consistent with an early, active control process associated with suppression, we found, within the first 500 ms, significant NT vs. PB decoding in 4-8 Hz theta activity over the frontal and posterior regions (Figure 3E, 3H). This significant decoding continued throughout the 3000 ms epoch. Theta power differences contributed to this decoding: we found that during the 200-400 ms window, retrieval suppression (vs. retrieval or passive viewing) led to enhanced midline and right prefrontal theta power (NT > T, *p*_corrected_ = .007, Figure 3I; NT > PB, *p*_corrected_ = .002, Figure S1G). After this early theta enhancement, suppression was associated with reduced theta power from 500 to 3000 ms (NT < T, theta: *p*_corrected_ = .004, NT < PB, theta: *p*_corrected_ < .001).

**Fig. 3.**
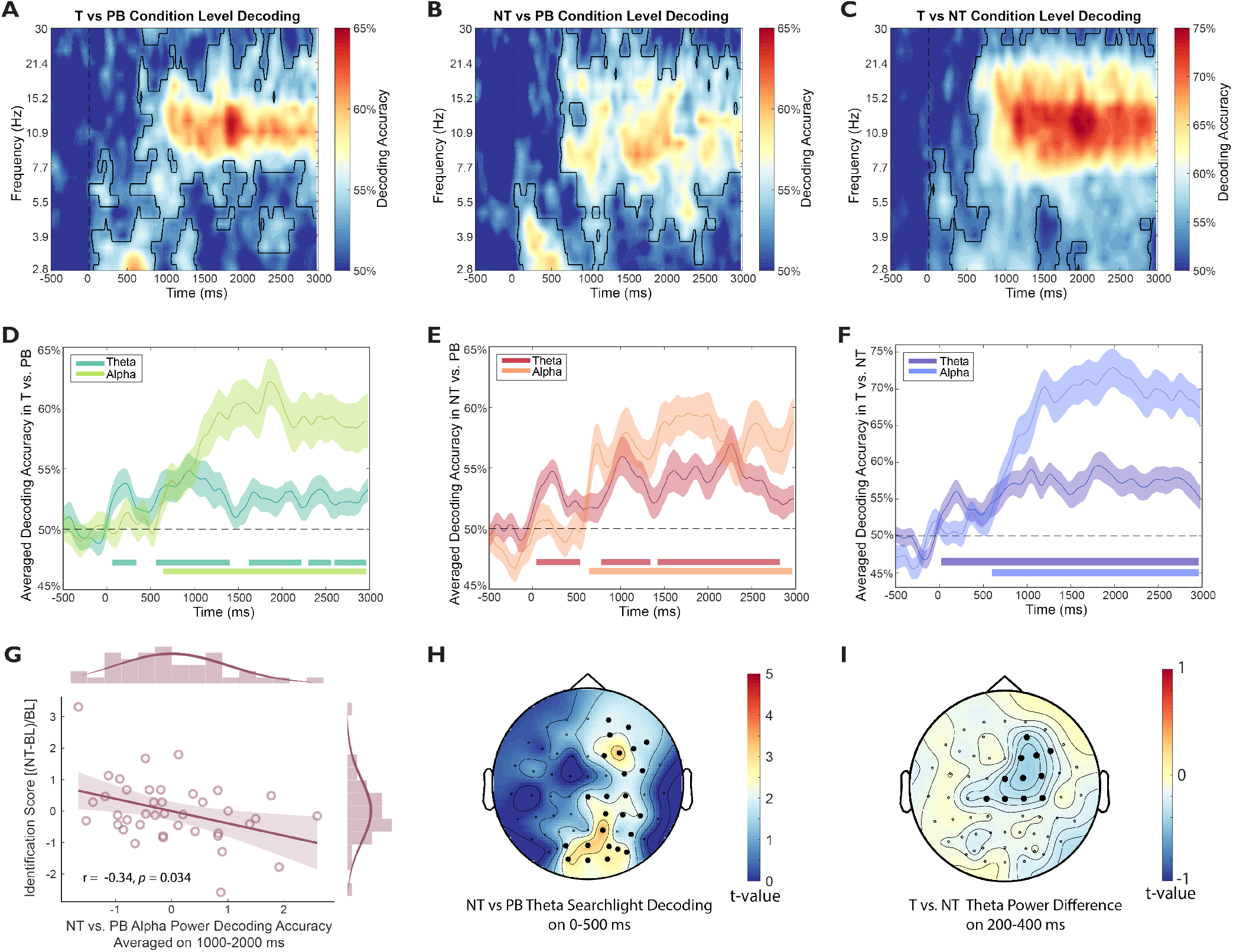
The Condition-Level Time-Frequency Domain Decoding. (A-C) Condition-level time-frequency decoding results. Frequency is log scaled with the colorbar denoting decoding accuracy. Black outlined highlight significant clusters against chance level (both cluster and permutation *α*s are set at 0.05). (D-F) Decoding accuracies in A-C are averaged on theta (4-8 Hz) and alpha (9-12 Hz) bands. Lines at the bottom denote significant clusters of averaged accuracy against chance level (50(G) The alpha-based No-Think vs. Perceptual Baseline decoding accuracies during 1,000-2,000 ms predicted later suppression-induced forgetting (i.e., higher decoding predicted a more negative score, or higher forgetting). (H) Theta power within 0-500 ms distinguished NT vs. PB over frontal and posterior brain regions in a channel searchlight decoding analysis. Significant electrodes were cluster corrected and are highlighted. (I) Theta power averaged from 200-400 ms was higher in NT than T. The increased theta power showed a frontal-central distribution. Significant electrodes were cluster corrected and are highlighted.

Retrieval suppression also could be distinguished based on alpha activity, and such effects were enduring. Indeed, 9-12 Hz alpha activity drove condition-level decoding performance between 500 to 3000 ms (Figure 3D-F) with retrieval suppression reducing alpha (NT <T, *p*_corrected_ < .001; NT<PB, *p*_corrected_ = .002, Figure S1A-F). Based on a recent study indicating that a 1000-2000 ms alpha power reduction may reflect reduced rehearsal during memory control (Fellner et al., 2020), we hypothesized that these alpha power reductions may have behavioral implications. Strikingly, during the same 1000-2000 ms as in prior research, the ability to decode NT versus PB based on alpha activity predicted suppression-induced forgetting on our *Identification* measure (r = -0.34, *p* =.034, Figure 3G, also see Figure S2B). This negative correlation suggests that reduced alpha power contributed to subsequent forgetting of suppressed content. In contrast, whereas alpha-based NT vs. PB decoding accuracies predicted suppression-induced forgetting, the ability to decode T from PB based on alpha power predicted retrieval-induced facilitation for Think items, with the difference of these two correlations being significant (*Detail*: z = 2.06, *p* =.039; Figure S2C). Together, these findings suggest that increases in early theta power and reductions in later theta/alpha power may be hallmarks of active suppression that make it qualitatively distinct from simply not-retrieving.

### Spatial Patterns in EEG Discern Individual Episodic Memories During Retrieval

Observing the suppression of individual memories requires an index sensitive to brain activity unique to each memory item so that the impact on suppression on that index may be tracked. We hypothesized that the spatiotemporal pattern of scalp-EEG as participants thought about each scene may contain information sufficient to distinguish that specific scene from all the others. To test this hypothesis, we performed a decoding analysis on scalp-EEG patterns during Think trials, during which participants actively reinstated associated scenes. Consistent with our hypothesis, time-domain EEGs distinguished between individual scene memories across the entire 0-3000 ms window (Figure 4A, *p*_corrected_ < .001). In sharp contrast, for Perceptual Baseline trials, above-chance decoding of individual items arose only in the 0-500 ms (to be precise, 60-640 ms, *p*_corrected_ < .001), but not in the 500-3000 ms window (Figure 4C). To directly compare item-level decoding between retrieval and PB, we repeated the analyses with 6 randomly sampled items from the Think condition, to match the item number in the Perceptual Baseline (see Methods). We found that Think trials showed higher item-level decoding accuracies than Perceptual Baseline trials during the 360-1180 ms (*p*_corrected_ < .001) and 1220-1540 ms window (*p*_corrected_ = .022, Figure 4K, purple lines).

**Fig. 4.**
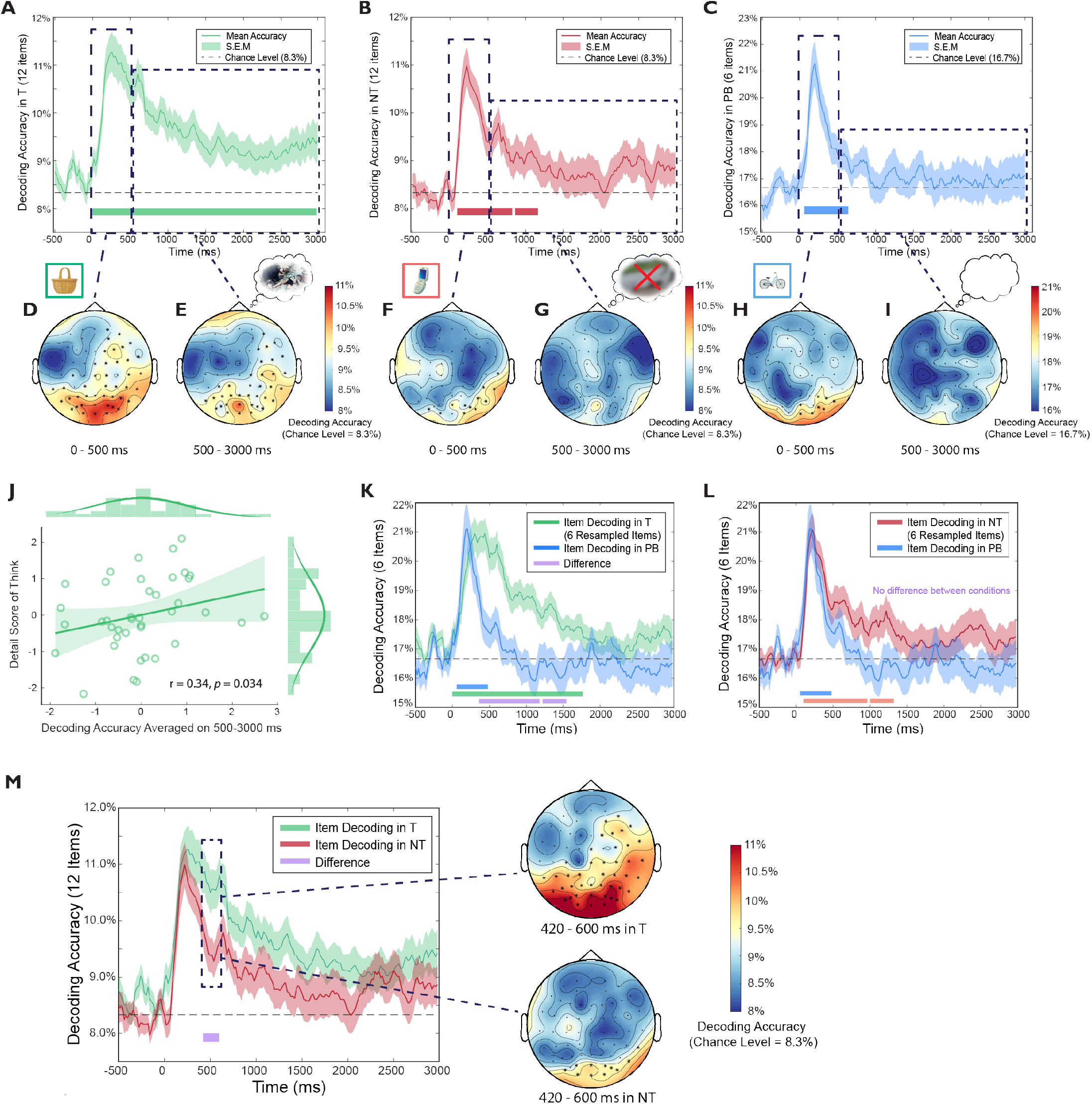
Item-level Time Domain Decoding. (A-C) The item-level decoding patterns (averaged across participants) in each retrieval condition. Lines at the bottom indicate significant time clusters against chance level, with permutation cluster correction (*α*s = 0.05). (D-I) Channel searchlight analyses of time domain decoding during an early (0-500 ms) and a later time window (500-3000 ms). The colorbar indicates decoding accuracy. Electrodes with significant decoding accuracies are highlighted (permutation cluster corrected, *α*s = 0.05). (J) During Think trials, decoding accuracies averaged on 500-3000 ms predicted the number of details recalled from emotional scenes. (K) Item-level decoding in the Think condition (using 6 resampled items) is higher than it is in the Perceptual Baseline condition from 360-1180 ms, *p*_corrected_ < .001 and from 1220-1540 ms, *p*_corrected_ = .022. Lines at the bottom indicate cluster-corrected significant time clusters against the chance level (green and blue for Think and Perceptual Baseline) or the difference between the two conditions (purple). (L) Item-level decoding in the No-Think condition (using 6 resampled items) is not significantly different from decoding in the Perceptual Baseline condition. Lines at the bottom indicate significant time clusters against the chance level (red and blue for No-Think and Perceptual Baseline conditions, respectively). (M) Retrieval suppression significantly reduced item-level decoding accuracies from 420-600 ms compared to retrieval (Think condition), with the right panel showing channel searchlight analyses on this time window.

Successful decoding of individual items in the early time window (0 – 360ms) likely reflects visual processing of unique object retrieval cues, which are present both for the object-scene pairs used in the Think condition, and in the single objects used in the PB condition. In the later 360-1540 ms time window, however, higher decoding during Think trials would need to be driven by an item-specific processing present in the Think condition but not in the PB condition. One possibility is that this later item-specific effect in the Think condition may reflect the reinstatement and maintenance of unique unpleasant scenes associated to the object cue, which may have gradually begun to emerge in awareness as they were recollected. Another possibility, however, is that item-level decoding in the Think condition may simply reflect more sustained attention to the unique object cues in that condition, relative to the PB condition, for which participants may have correctly concluded that retrieval was unnecessary.

To distinguish these possibilities, we examined brain regions giving rise to above-chance decoding during Think trials using searchlight decoding (see Methods). If greater decoding of individual items in the Think condition reflected sustained attention on object cues, successful decoding may be restricted to visual processing regions involved in object perception. Indeed, during the first 500 ms, occipital EEGs primarily drove the significant decoding in general, consistent with a primary role of visual-perceptual cue processing (Figure 4D). In contrast, during the latter 500-3000 ms interval, significant decoding rested on a distributed set of regions implicated in memory retrieval such as the right prefrontal and parietal-occipital cortex (Figure 4E). This finding suggests that item-level decoding beyond the first 500 ms is not dominated by object cue attention, but rather by the reinstatement of the associated scene memories. Converging with this possibility, item-level decoding performance during the latter 500-3000 ms time window predicted later performance on the Detail measure of scene memory (r = 0.34, *p* = .034, Figure 4J), whereas decoding during the early 0-500 ms time window did not (r = 0.01, *p* = .946).

Unlike during Think trials, the same searchlight analysis during Perceptual Baseline trials showed that significant decoding in the 0-500ms window arose over a small cluster of occipital electrodes. The restriction of decoding success to occipital cortex suggests that classification hinged on visual object processing during that period (Figure 4H). After this initial window, the latter part of the trial from 500-3000 ms showed no significant decoding at any electrode (Figure 4I; similar searchlight results were obtained when using 0-360 and 360-1540 ms windows, see Figure S3A).

In sum, during retrieval, time-resolved EEG patterns suggest a staged cued-recall process: during the 0-500 ms window, EEG patterns could discern perceived items over occipital regions; during 500-3000 ms, EEG patterns could distinguish among retrieved items over fronto-parietal-occipital regions. Furthermore, higher item-level decoding accuracies predicted better scene memory only in this latter, 500-3000 ms time window.

### Suppressing Retrieval Weakens and Abolishes Item-specific Cortical Patterns

Having established that the retrieval of individual scene memories can be indexed and tracked, we next sought to use this index to determine how and when suppression affected cortical patterns relating to individual memories. We therefore examined whether retrieval suppression modulated item-specific cortical EEG patterns.

We hypothesized that item-level decoding during No-Think trials would be possible initially, as participants focused their attention on the visually unique reminder cues, but that suppression would limit successful decoding throughout the remainder of the trial. Indeed, in the No-Think condition, item-level decoding accuracy was above chance initially, and remained so until 1160 ms (*p*s_corrected_ < .028); decoding accuracy then dropped to chance-levels for the remainder of the 3000ms trial. Consistent with the Think and Perceptual Baseline analyses, we used a priori defined time windows from 0-500 and 500-3000 ms to characterize the EEG scalp distributions contributing to decoding success. During the 0-500 ms window, item-level decoding was driven by occipital activity, resembling the EEG distributions found in the Perceptual Baseline condition during the same window (Figure 4F, 4H). Strikingly, during the 500-3000 ms, there were no brain regions that contributed significantly to item-level decoding (Figure 4G), suggesting that suppression had abolished evidence for cortical reinstatement of scene memories.

In addition to scalp EEG distributions revealed by the channel searchlight analysis, confusion matrices of item-level decoding provided converging evidence supporting the hypothesized stages of retrieval suppression: we observed significant above-chance item-specific classifications in all three conditions during the first 500 ms, when cue-processing might be expected to predominate; in contrast, distinctive classification patterns remained only in the Think condition during 500-3000 ms (Figure S3C-E). Thus, suppression reduced cortical patterns during No-Think trials to the extent that they were as uninformative as items in our perceptual baseline condition, in which no scene retrieval was possible.

To precisely characterize of the temporal dynamics of retrieval suppression, we contrasted the time-dependent evolution of item-specific cortical patterns between retrieval suppression and both the retrieval and perceptual baseline conditions. A direct comparison of Think vs. No-Think item-level decoding revealed that retrieval suppression reduced decoding accuracies from 420 to 600 ms (*p*_corrected_ = .044, Figure 4M left panel). Searchlight analyses during 420-600 ms revealed that, whereas voluntary retrieval engaged item-specific brain activity over frontal-parietal-occipital regions, retrieval suppression was only associated with occipital activity (Figure 4M right panel). When No-Think trials were directly compared to Perceptual Baseline trials (using 6 randomly sampled items from the No-Think condition), there were no significant decoding accuracy differences during the entire 0-3000 epoch (none of the differences survived permutation correction, see Figure 4L).

Reduced decoding accuracy for individual No-Think items in the 420-600ms window suggests that the retrieval stopping process may begin to exert its first effects within this window, a possibility consistent with findings from our condition-level decoding analyses. We next sought to determine whether prefrontal-control processes were linked to suppressed item-level decoding. Consistent with this possibility, we found that in the No-Think (vs. Think) trials, reduced item-level decoding was preceded by enhanced 200-400 ms theta power over midline and right prefrontal cortex (Figure 3I). Critically, theta power elevation across this region positively correlated with the 420-600 ms decoding accuracy reduction (r = 0.30, *p* = .064, Figure S3F), suggesting that processes indexed by higher theta power (No-Think > Think) contributed to lower item-specific decoding accuracies (No-Think < Think).

Together with the evidence for suppression-specific patterns in the condition level analysis, these item-level decoding results reveal a precise timeline of how retrieval suppression unfolded: inhibitory control was engaged within the first 500 ms upon encountering a unwelcome reminder cue, presumably before the cue-to-memory conversion process completed, to obstruct retrieval and prevent reinstatement from happening. This early control weakened, and eventually abolished memory-specific cortical patterns during 500-3000 ms.

### Rapid and Sustained Suppression of Individual Memories Led to Their Forgetting

To understand how the temporal dynamics of retrieval suppression influenced later forgetting of suppressed content, we divided participants into *High-* vs. *Low-Suppression Groups* based on a median-split of suppression-induced forgetting scores. We focused on below-baseline forgetting (i.e., NT-minus-BL *Detail* scores) using our detail measure of scene recall. We tested the hypothesis that successful suppression-induced forgetting was associated with a greater reduction in decoding accuracy during No-Think trials compared to Think trials, compared to unsuccessful forgetting. In the High-Suppression group (Figure 5A), suppression significantly reduced item-specific decoding accuracy during No-Think (vs. Think) trials during two time windows: 300-680 ms (*p*_corrected_ = .006) and 1140-1400 ms (*p*_corrected_ = .031). By contrast, in the Low-Suppression group (Figure 5B), the same comparison revealed no NT vs. T decoding accuracy differences, indicating that evidence for item-specific decoding remained possible for this group, despite their efforts to suppress. In the high forgetting group, the observed differences may reflect an early disruption of cue-to-memory conversion processes occurring at around 500 ms, and a later weakening of item-specific cortical reinstatement between 1000-1500 ms. Corroborating a role of early and timely suppression in forgetting, item-level decoding accuracy during the early 300-680 ms window predicted later suppression-induced forgetting across all participants (r = 0.35, *p* = .027, Figure 5C). Thus, the more effectively participants suppressed unwanted memories during the 300-680 ms window, the more successful was the later forgetting of scene details.

**Fig. 5.**
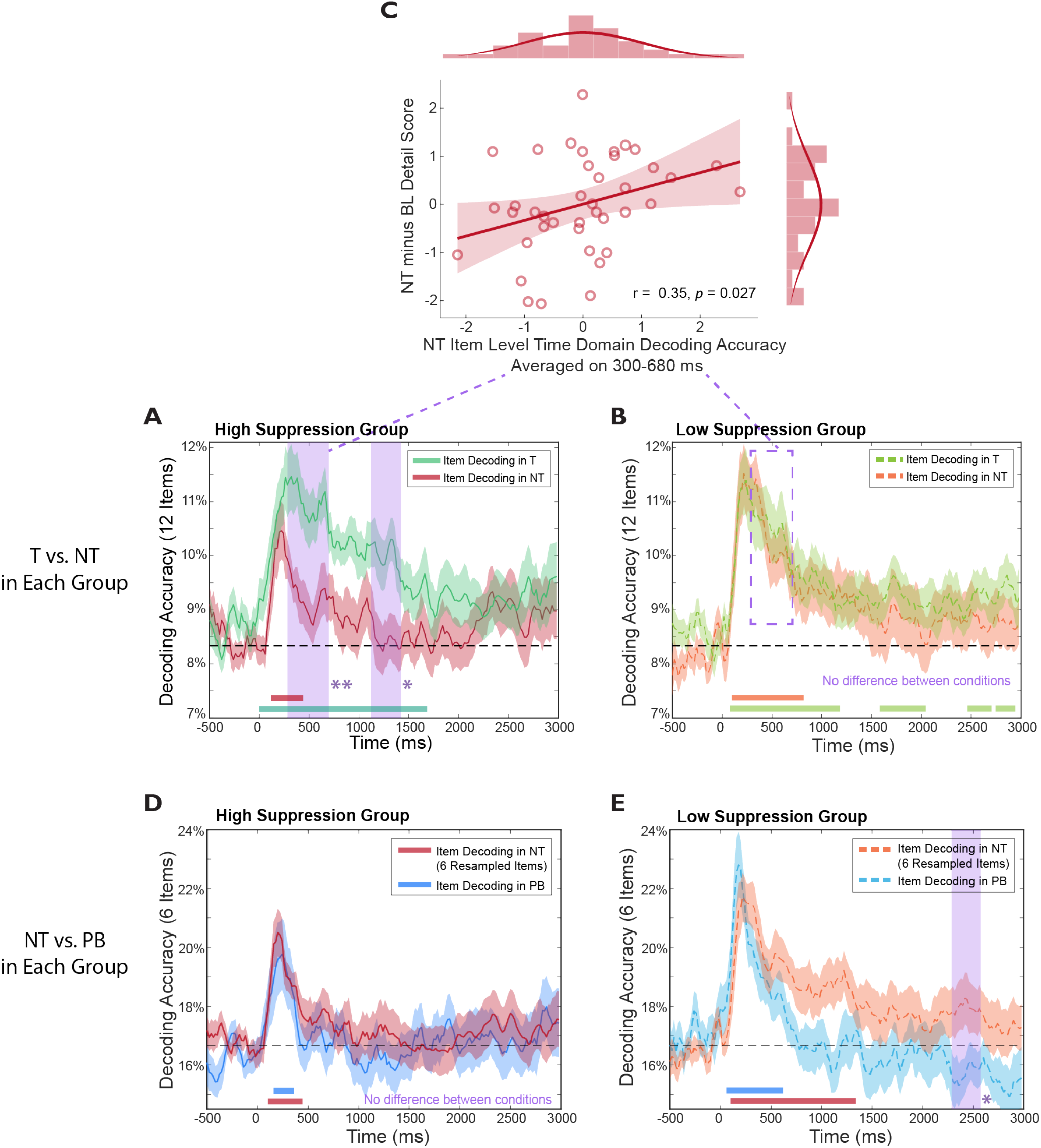
Item-level Decoding Results in Highand Low-Suppression Groups. (A, B) Comparisons between Think and No-Think item-level decoding in High-/Low-Suppression Groups, respectively. In the High-Suppression Group, the Think vs. No-Think difference was significant during the 300-680 ms and 1140-1400 ms windows, whereas no differences were found in the Low-Suppression Group. (C) Across both groups, the averaged decoding accuracy during the 300-680 ms. window positively correlated with participant’s suppression-induced forgetting (i.e. No-Think Baseline of the detail index). (D, E) Resampled item-level decoding comparisons between the No-Think and Perceptual Baseline conditions in the Highand Low-Suppression Groups, respectively. In the High-Suppression Group, the No-Think condition did not differ from the Perceptual Baseline condition in item-level decoding accuracy, despite both showing above chance decoding within the 0-500 ms window. In the Low-Suppression Group, in contrast, a significant difference between the No-Think and the Perceptual Baseline conditions was observed during the 2300-2560 ms window. Colored bars at the bottom of each figure denote time clusters that were significantly above chance (permutation corrected, one-sided *α*s = 0.05). Purple dashed outlines denote significant time clusters between conditions/groups (permutation corrected, two-sided *α*s = 0.05).

We next compared item-level decoding accuracy between the No-Think (using 6 randomly sampled items) and Perceptual Baseline conditions in the Low and High-Suppressor groups. Strikingly, we found no between-condition differences in the *High-Suppression Group* (Figure 5D), indicating that suppression reduced pattern information so effectively that the brain activity contained no evident item-specific content, mimicking those trials in which there was actually no scene to reinstate. In contrast, participants from the *Low-Suppression Group* showed significantly higher decoding accuracies during No-Think trials compared to Perceptual Baseline trials, primarily toward the end of the suppression epoch (i.e., 2300-2560 ms, *p*_corrected_ = .029, Figure 5E, purple dashed outline). Thus, less successful forgetting was associated with relapses in the activation of suppressed content during sustained control of unwanted memories. Together, these results highlight that not only early and rapid, but also sustained control are important in successful suppression-induced forgetting.

### Theta and Alpha Oscillations Track Item-Level Perception and Reinstatement Processes, Respectively

We sought converging evidence for the active suppression of individual memories by tracking item-specific oscillatory activity in the theta and alpha bands. Theta and alpha activity have been implicated in perceptual and memory-related processes, such that theta may reflect sensory intake and hippocampo-cortical communication loops (Bastos et al., 2015; Colgin, 2013), and alpha may track neocortex-dependent memory reinstatement processes (Staresina et al., 2019; Staresina & Wimber, 2019). If so, posterior theta activity may enable item-specific decoding of the cue objects themselves, whereas alpha activity may enable decoding of reinstated scenes.

In all three conditions, we found that theta activity in the 0-500ms window over occipital regions significantly distinguished among individual items, consistent with theta’s putative role in visual processing of individual cue objects (*p*s_corrected_ < .001, Figure 6A-C, also see Figure 6D-G). During the 500-3000 ms window in which scene recollection could unfold, both theta and alpha power drove significant decoding accuracy during Think trials (theta: *p*s_corrected_ < .027; alpha: *p*s_corrected_ < .039, Figure 6D). Critically, however, retrieval suppression during No-Think trials abolished any evidence for item-specific decoding based on theta or alpha band activity (Figure 6E). There was short-lived theta-driven decoding in Perceptual Baseline trials, which may reflect occasional perceptual processing of objects (theta: *p*s_corrected_ < .011, Figure 6F). Channel searchlight analyses during the 500-3000 ms window revealed that alpha activity over the occipital-parietal region contributed to decoding performance in the Think condition, but did not in either the No-Think or Perceptual Baseline conditions (see Figure 6H). These findings support the possibility that alpha activity is linked with scene-specific memory reinstatement processes and not simply to object perception. If so, the lack of significant alpha-based decoding in No-Think trials reflects the abolition of memory reinstatement processes arising due to active suppression.

**Fig. 6.**
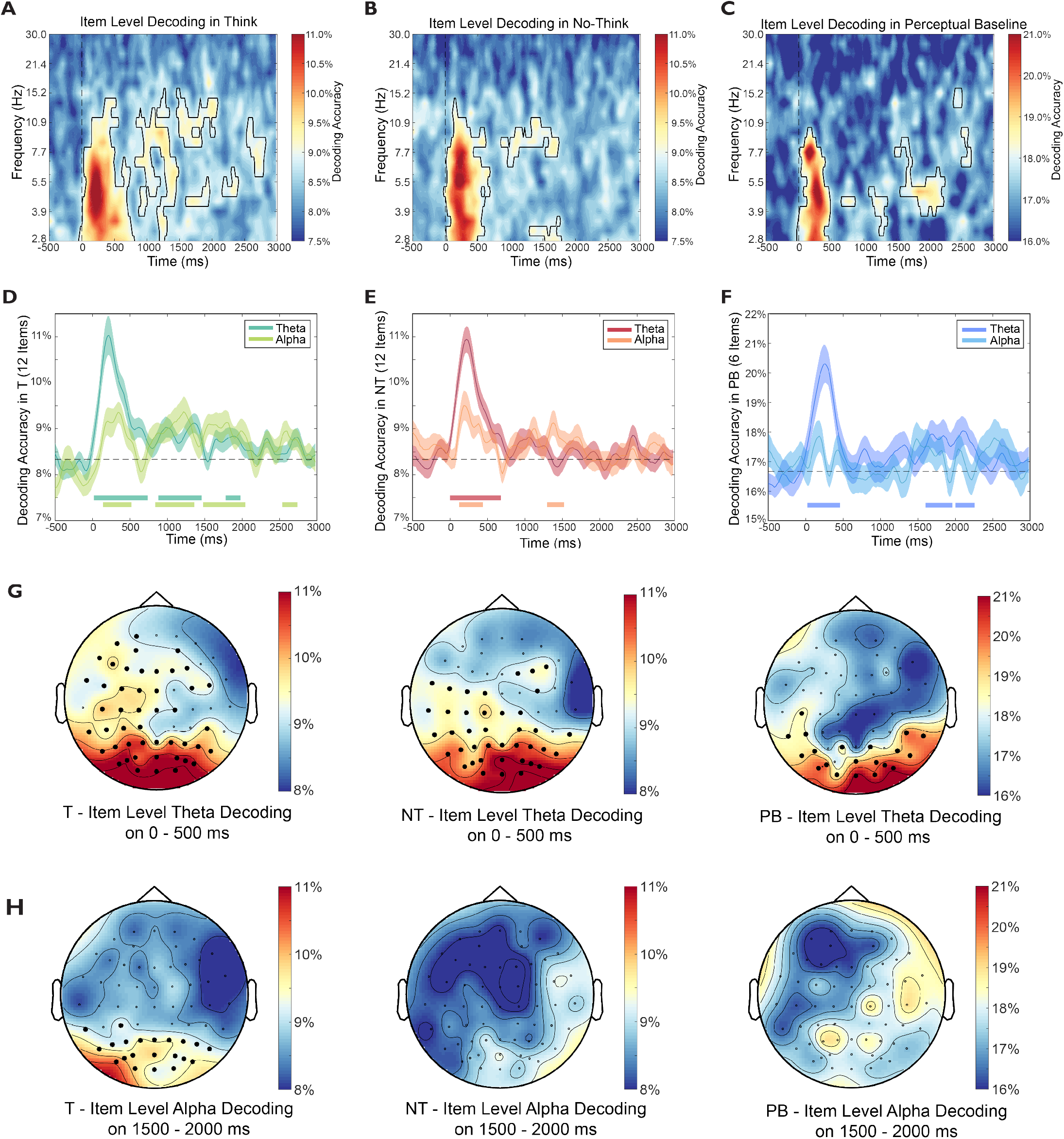
Item-level Time-Frequency Domain Decoding. (A-C) Item-level time-frequency decoding results. Frequency is log scaled and the colorbar denotes decoding accuracy. The black outline highlights significant clusters against chance levels (both cluster *α* and permutation *α* are 0.05, one-sided). (D-F) Decoding accuracies in A-C are averaged on theta and alpha bands. Horizontal bars denote significant clusters of the band-averaged accuracies against chance level (cluster corrected, one-sided *α*s = 0.05). (G) Item-level theta searchlight during the 0-500 ms window showed an occipital distribution in all three conditions. Significant channels are highlighted (permutation cluster corrected with one-sided *α*s = 0.05). (H) Item-level alpha searchlight during the 1500-2000 ms window showed that only in the Think condition was alpha power able to distinguish among items. The alpha searchlight decoding in the Think condition originated from the posterior region. Significant channels are highlighted (permutation cluster corrected with one-sided *α*s = 0.05).

## Discussion

Suppressing memory retrieval requires effort; it is not simply neglecting to engage retrieval when an unwelcome reminder appears, but instead involves an active inhibition process (Anderson & Hulbert, 2020; Wimber et al., 2015). Applying multivariate pattern analyses during the think/no-think task, we observed, for the first time, how individual aversive memories are suppressed in real time. Our precise chronometry of retrieval suppression provides new knowledge about the time windows and neural activity critical to achieving successful forgetting. We found that effective forgetting is associated with 1) the rapid deployment of inhibitory control in suppressing cortical patterns within the first 500 ms, supported by enhanced midfrontal theta activity during efforts to stop retrieval; and 2) sustained control applied to abolish item-specific cortical EEG patterns reflected in the spatial pattern of theta and alpha activity during the 500-3000 ms window.

Three findings suggest that an early, active control process truncates retrieval of highly specific, individual memories, inducing later forgetting. First, when a reminder cue appeared, within 500 ms retrieval suppression enhanced midfrontal and right prefrontal theta activity relative to active retrieval and also relative to a perceptual baseline condition in which scene retrieval was impossible. Given evidence linking frontal midline theta and inhibitory control (Cavanagh & Frank, 2014; Crespo-García et al., 2021; Nigbur et al., 2011), this finding is consistent with the possibility that attempts to stop the retrieval process engaged inhibitory control. This finding suggests a rapid onset of inhibitory control in the face of an unwelcome reminder, but does not, by itself, link that control process to the successful exclusion of unwanted memories from awareness.

Second, whereas we detected significant item-specific brain activity during active retrieval, retrieval suppression reduced the ability to detect individual items during the 420-600 ms time window. The ability to detect reduced item-specific activity in such an early time window indicates that suppression rapidly interrupts the retrieval process. Estimates based on intracranial recordings suggest that beginning at around 500 ms, hippocampus-dependent pattern completion would normally trigger cortical reinstatement of target memories, accompanied by vivid recollection (Colgin, 2016; Lavenex & Amaral, 2000; Staresina & Wimber, 2019). Given this timing, successful retrieval suppression ideally should target prior to this time window to pre-empt or truncate the cue-to-memory conversion processes, preventing memories from being reinstated. Indeed, our putative index of inhibitory control predicted reduced item-specific EEG activity: we found that elevated theta power in the 200-400 ms window predicted later reductions of item-level decoding accuracy in the 420-600ms window. These findings suggest that enhanced inhibitory control disrupted the cue-to-memory conversion process to prevent aversive memories from being retrieved, but it does not link such changes in cortical reinstatement to later forgetting of the suppressed content.

Third, we found that reduced item-specific cortical pattern information during this early time window predicted later suppression-induced forgetting. Specifically, whereas those participants showing high suppression-induced forgetting exhibited significantly reduced item-level decoding accuracies during suppression, compared to retrieval in the 300-680 ms window, Low-Suppression participants did not. In general, across all participants, reduced No-Think item decoding accuracies within the 300-680 ms window predicted later suppression-induced forgetting. These results link the early engagement of control not only to reduced reinstatement, but also to an increased capacity to forget the suppressed content. Given that hippocampus-dependent pattern completion processes emerge at around 500 ms (Staresina & Wimber, 2019), this finding again suggests that for successful forgetting to occur, top-down inhibitory control should be engaged quickly before and during the cue-to-memory conversion time window, preventing cortical reinstatement.

The temporal evolution of item-specific cortical patterns suggests that whereas rapid control is important to successful forgetting, sustained control also is necessary. Whereas retrieval suppression weakened the ability to detect item-specific cortical patterns starting from ∼400 ms after cue-onset, individual memories could still be identified until 1200 ms post-cue. Residual item-specific cortical patterns during the 420-1200 ms window clearly call for sustained control to ensure that unwanted memories are suppressed. The ability to detect item-specific cortical patterns was fully abolished by 1200 ms for the remainder of the 3000ms trial. The maintenance of control over this longer time period appears to be reflected in reduced alpha power throughout the trial. Together, these temporal characteristics reveal a timeline for the suppression of aversive scenes: early control processes truncate retrieval during the perception-to-memory conversion time window (e.g., ∼420-600 ms), with sustained control processes down-regulating unwanted memories (e.g., ∼1200 ms), eventually abolishing item-specific cortical patterns (1200-3000 ms).

Two additional findings underscore the importance for sustained control in the successful forgetting of unwanted memories. Although early control clearly was instrumental to successful forgetting, we also found evidence that activity in later time windows was also functionally relevant. First, those participants showing higher suppression-induced forgetting showed significantly reduced item-level decoding accuracies during suppression than during retrieval in the 1140-1400 ms time window, suggesting the functional importance of sustained control. Second, low-suppression participants showed evidence of an ironic rebound effect later in the trial: retrieval suppression was associated with significantly higher decoding accuracies than our perceptual baseline trials in the 2600-2800 ms time window. This apparent rebound effect in cortical reinstatement suggests that participants who later showed less successful forgetting suffered relapses in controlling unwanted memories, particularly towards the end of retrieval suppression (van Schie & Anderson, 2017). Taken together, these two findings illustrate that successful forgetting requires sustained suppression of individual memories during the prolonged cortical reinstatement time window.

Our item-level decoding results during voluntary retrieval trials (i.e., Think trials) provide converging evidence for our staged view of how cued memory recall unfolds. To determine whether sustained item-level decoding during Think trials might simply reflect persisting attention to individual object cues, we showed that 1) the early (0-500 ms) vs. late (500-3000 ms) decoding patterns were characterized by distinct EEG spatial distributions, and 2) only the 500-3000 ms item-level decoding accuracy predicted more detail of scene recall of Think items on the later test. These results suggest that whereas the early decoding pattern reflects perceptual processes acting on item-specific cues, the later decoding pattern likely reflects the successful recollection of the accompanying scene. Consistent with this interpretation, both theta and alpha power contributed to item-level decoding during voluntary retrieval, with an early onset of occipital theta activity followed by parietal-occipital alpha activity. Theta and alpha activities have been implicated in perceptual and memory-related processes, such that theta may reflect sensory intake and hippocampo-cortical communication loops (Bastos et al., 2015; Colgin, 2013). Relatedly, linking behavioral oscillation and neural oscillation, a recent study demonstrated a prominent role of theta rhythm in memory retrieval (Ter Wal et al., 2021). Regarding alpha, previous research suggests that alpha may track neocortex-dependent memory reinstatement processes (Staresina et al., 2019; Staresina & Wimber, 2019) Decoding patterns during Perceptual Baseline trials provided converging support for this account: when participants viewed object cues that lacked any associated scene memory, only occipital theta activity in the 0-500ms window drove significant item-level decoding, ruling out any contribution of scene retrieval.

If the foregoing staged view of retrieval is correct, then item-specific decoding based on alpha-band activity after initial cue processing may reflect the reinstatement of individual scenes. Indeed, previous research has found that memory reinstatements are associated with alpha oscillations. For example, in a directed forgetting task, Fellner et al. (2020) reported alpha power increases 1000-2000 ms following to-be-remembered cues, which were associated with selective rehearsal (see also Bäuml et al., 2008; Hanslmayr et al., 2012; Xie et al., 2020). Mirroring this, we found that voluntary retrieval enhanced alpha power during the same 1000-2000 ms window when reinstatement of the associated scene would be expected (Figure S1H-M). If this interpretation is correct, then the reduced alpha power relative to our perceptual baseline condition (and alpha-based item-level decoding performance), likely reflects the outcome of suppressing scene reinstatement. Critically, higher decoding based on alpha activity during retrieval suppression, relative to the perceptual baseline condition predicted later suppression-induced forgetting. Suppression-induced alpha power reductions may reflect reduced memory reinstatement (Hanslmayr et al., 2012; Waldhauser et al., 2015), which contributed to episodic forgetting.

Taken together, our findings show that for successful retrieval suppression and forgetting, inhibitory control needs to be both fast and sustained. On the one hand, early enhanced frontal theta disrupted cue-to-memory conversion, truncating the reinstatement of individual aversive scene memories within the first 500 ms upon seeing the cues. On the other hand, sustained control weakened and eventually abolished item-specific cortical EEG patterns during the 500-3000 ms time window, reflected in reduced alpha activity. In contrast, both diminished early control and relapses during later sustained control compromised successful voluntary forgetting of suppressed content. By tracking the precise timing and neural dynamics of retrieval suppression in modulating individual memories, our results may inform future research on when and how to intervene during retrieval suppression to improve people’s ability to forget unwanted memories.

## Materials and Methods

### Experimental Subject Details

Fourty-one participants (mean age = 19.57, age range: 18-23 years, 26 females) were recruited from the University of Hong Kong. One participant was excluded due to non-compliance of task instructions (details see Materials and Procedure). Ethical approval was obtained from the Human Research Ethics Committee of The University of Hong Kong.

### Materials and Procedure

We used 42 object-scene picture pairs from Küpper et al. (2014). Scenes depict aversive contents such as natural disasters, assault, injury, etc. Each object resembled an item from its paired negative scene, thus establishing naturalistic and strong associations. Six pairs were used for instruction and practice purposes. The remaining 36 pairs were equally divided into 3 sets, with 12 pairs in each of three following conditions: Think, No-Think, and Baseline. Picture pairs used in the three conditions were matched on valence and arousal, and were counterbalanced across participants. Another 6 objects without any paired scenes were used as Perceptual Baseline trials, which did not involve any memory retrieval. Participants completed the following sessions in order: Encoding, Think/No-think (TNT) and Cued Recall. At the end of the study, participants completed a 3-item, instruction compliance questionnaire.

#### Encoding

Participants were presented with 42 object-scene pairings, plus 6 objects from Perceptual Baseline. Each object-scene pair was presented on an LCD monitor for 6 s with an inter-trial-interval (ITI) of 1s. Participants were instructed to pay attention to all the details of each scene, and to associate the left-sided object and the right-sided scene. They next completed a test-feedback session, in which each object was presented up to 4 s until participants pressed a button indicating whether they could recall the associated scene or not. If participants gave a ‘yes’ response, they were presented with three scenes from the learning phase and needed to identify the correct one within another 4 s. Regardless of accuracy, the correct pairing would be presented for 2.5 s. This test-feedback cycle repeated until participants reached 60% accuracy. Twenty-six participants reached this criterion in the first cycle, 13 participants in two, and 1 in three. Following the test-feedback cycles, participants completed a recognition-without-feedback test, so as to confirm that items from different conditions were encoded at comparable levels before the TNT session (*p*s >.104).

#### TNT

Participants were presented with 24 objects from the 36 object-scene pairings, with 12 objects in each of the Think or No-think conditions, respectively. The remaining 12 objects were not shown in the TNT and were used in the Baseline condition. These 24 objects were presented in either yellow- or blue-colored frames indicating think and no-think conditions, with colors counterbalanced across participants. Six objects (without any pairing scenes) were presented in white-colored frames and served as Perceptual Baseline trials. Thus, 30 unique objects were shown in the TNT session. Each object was presented 10 times, resulting in a total of 300 trials. Each trial began with a fixation cross (2-3s), followed by the object in a colored frame for 3s. The ITI was 1 s.

For Think trials, participants were instructed to try their best to think about the objects’ associated scenes in detail, and to keep the scenes in mind while the objects remained on the monitor. For No-Think trials, participants were given direct-suppression instructions: they were told to pay full attention to the objects while refraining from thinking about anything. If any thoughts or memories other than the objects came to mind, they needed to try their best to push the intruding thoughts/memories out of their mind and re-focus on the objects. Participants were also prohibited from using any thought substitution strategies (i.e., thinking about a different scene). For Perceptual Baseline trials, participants were simply instructed to focus on the object.

#### Cued Recall

Following the TNT session, participants were presented with each of the 36 objects from Think, No-Think and Baseline conditions. Each object was presented at the center of the monitor, alongside a beep sound prompting participants to verbally describe the associated scenes within 15 s. The ITI was 3 s. Participants’ verbal descriptions were recorded for later scoring. Perceptual Baseline objects were not shown in this recall test because they were not paired up with any scenes.

#### Cued Recall Analyses

Two trained raters who were blind to experimental conditions coded each of the verbal descriptions along three dimensions following the criteria used in in a previous study (Küpper et al., 2014), namely *Identification, Gist* and *Detail*. Each measure focused on different aspects of memories: Identification referred to whether the verbal description was clear enough to correctly identify the unique scene, and was scored as 1 or 0. Inconsistent ratings were resolved by averaging 0 and 1, resulting in a score of 0.5. Gist measured whether participants’ verbal descriptions contained critical elements pertaining to the scene’s main themes. Two independent raters identified two to four gists for each scene (Küpper et al., 2014). We scored gist as proportion, using the number of correct gists from participants’ verbal report divided by all possible gists for each scene. Detail measured how many correct meaningful segments were provided during the verbal description, and was scored on the number of details. Interrater agreement for the scoring of all three measures was high: Identification r = 0.71, Gist r = 0.90, Detail r = 0.86.

#### EEG Recording and Preprocessing

Continuous EEGs were recorded during the TNT session using ANT Neuro eego with a 500 Hz sampling rate (ANT, The Netherlands), from 64-channel ANT Neuro Waveguard caps with electrodes positioned according to the 10-5 system. The AFz served as the ground and CPz was used as the online reference. Electrode impedances were kept below 20 kilo-ohms before recording. Eye movements were monitored through EOG channels.

Raw EEG data were preprocessed in MATLAB using EEGlab Toolbox (Delorme & Makeig, 2004) and ERPlab Toolbox (Lopez-Calderon & Luck, 2014): data were first downsampled to 250 Hz, and were band-passed from 0.1 to 60 Hz, followed by a notch filter of 50Hz to remove line noise. Bad channels were identified via visual inspection, and were removed and interpolated before re-referencing to common averages. Continuous EEG data were segmented into -1000 to 3500 ms epochs relative to the cue onset, and baseline corrected using -500 to 0 ms as baseline period. Next, independent component analyses (ICAs) were implemented to remove eye blinks and muscle artifacts. Epochs with remaining artifacts (exceeding ± 100 µV) were rejected. The numbers of accepted epochs used in all following analyses were comparable across Think (Mean ± SD, 100.33 ± 11.57) and No-think (103.18 ± 10.61) conditions. Valid trials number in Perceptual Baseline is 56.58 ± 3.23. All EEG analyses were based on 61 electrodes, excluding EOG, M1, M2, AFz (ground) and CPz (online reference).

#### Condition-/Item-level Decoding with Time Domain EEG

Decoding analyses were conducted in MATLAB using scripts adapted from (Bae & Luck, 2018), which used a support vector machine (SVM) and error-correcting output codes (ECOC). The ECOC model combined results from several binary classifiers for prediction output in multiclass classification.

In condition-level decoding, we used one-vs-one SVMs to perform pairwise decoding among the three conditions (Think vs. Perceptual Baseline, No-Think vs. Perceptual Baseline, and Think vs. No-Think). For Think vs. Perceptual Baseline and No-Think vs. Perceptual Baseline condition-level decoding, we first subsampled trials in T/NT to be comparable with Perceptual Baseline so that each condition had about 56 trials. We next divided EEG trials from each condition into 3 equal sets and averaged EEG epochs within each set into sub-ERPs to improve signal-to-noise ratio. The decoding was achieved within each participant from -500 to 3000 ms using these sub-ERPs in a 3-fold cross validation: each time 2 of the 3 sub-ERPs are used as training dataset with the condition labels, and the remaining one was used as testing dataset. After splitting training and testing datasets, sub-ERPs were both normalized using the mean and standard deviation of training dataset to remove ERP-related activity. This process was conducted on every 20 ms time point (subsampled to 50 Hz), and repeated for 10 iterations. We were comparing condition-level decoding accuracy against its chance level, 50%, given two conditions were involved in each pairwise decoding.

For item-level decoding, we used one-vs-all SVMs to decode each individual stimulus within each condition, separately. Decoding procedures were the same as condition-level decoding. Thus, the trial numbers of each stimulus are first matched to the least one within each participant (at most 10 trials, if no trial was rejected). Then, all trials of each stimulus were divided into 3 sets before averaging and the 3-fold cross validation. Both training dataset and testing dataset were normalized using the mean and standard deviation of training dataset. The decoding process was conducted on every 20 ms time point and for 10 iterations (results remained the same for up to 100 iterations, see supplementary Figure S3G). For Think and No-Think conditions, the chance levels were 1/12 (8.33%) given that there were 12 unique stimuli in each of these two conditions. For Perceptual Baseline trials, the chance level was 1/6 (16.67%).

Given we had different item numbers in Perceptual Baseline (6 items) and Think/No-Think (12 items), in order to directly compare the decoding accuracy in Think or No-Think with Perceptual Baseline, we conducted a resampled decoding in Think and No-Think, respectively. The resampled decoding is similar to the normal decoding, except that during each iteration we randomly selected 6 out of all 12 items before dividing and averaging into 3 sets. Considering the randomization used only half of the items, we increased iterations to 20 times. An item-level decoding with 20-iterations was also rerun in Perceptual Baseline, to be compared with the resampled decoding.

#### Condition-/Item-level Decoding with Time-Frequency Domain EEG

Time domain EEG was wavelet transformed into timefrequency domain data in Fieldtrip Toolbox (Oostenveld et al., 2011) before decoding. Frequencies of interest increased logarithmically from 2.8 Hz to 30 Hz, resulting in 22 frequency bins. Wavelet cycles increased linearly along with frequencies from 3 to 7. Then the decoding was conducted for each frequency bin data across time in the same procedure as described in Condition-/Item-level Decoding with Time Domain EEG (as if treating each frequency bin data as a time domain data).

#### Channel Searchlight Decoding

Both condition- and item-level decoding used EEGs from all 61 channels as features. To examine which electrodes contributed the most to the decoding accuracy, we conducted a channel searchlight decoding using subsets of the 61 channels as features (Treder, 2020).

Specifically, we first divided all channels into 61 neighbourhoods, centering each channel according to its location (conducted in Fieldtrip Toolbox (Oostenveld et al., 2011) via ft_prepare_neighbours() function using ‘triangulation’ method). Immediately neighbouring channels were clustered together, resulting in 6.39±1.50 channel neighbours for each channel (with overlaps). Then the time domain EEG was averaged on time windows of interest, i.e., averaged on 0-500 ms, 500-3,000 ms, etc., to inspect the decoding topographical distribution on different time windows. The rest of the procedure was the same as time domain EEG decoding: we divided data into 3 sets and averaged within each set before splitting training and testing datasets; then we normalized them using mean and standard deviation of training sets. Finally, the decoding was conducted with a 3-fold cross validation and 10 iterations. Theta/alpha searchlight was conducted in the same way as time-domain searchlight, after averaging timefrequency power on respective oscillation range (theta: 4-8 Hz; alpha: 9-12 Hz).

#### Time Frequency Analyses

Six electrode clusters were selected for Time Frequency analyses: left parietal (CP3/5, P3/5), parietal (Pz, CP1/2, P1/2), right parietal (CP2/4, P2/4), frontocentral (FC1/2, C1/2, FCz, Cz), left prefrontal (AF3, F3/5) and right prefrontal (AF4, F4/6).

Time frequency transformation was performed using the same parameters as in decoding analyses in Fieldtrip (Oostenveld et al., 2011), with additional decibel baseline normalization using power on -500 to -200 ms. We focus on the early theta power change on 200-400 ms which is indicator of inhibitory control (Cavanagh & Frank, 2014; Nigbur et al., 2011), and theta and alpha power change on a post hoc late time window (500-3000 ms) following condition level decoding results.

#### Correlation Analyses

We calculated Spearman’s Rho for all correlations. In condition-level decoding, memory of Think and No-think was normalized by subtracting and then divided by Baseline memory, then correlated with time domain condition-level decoding accuracy on 500-3000 ms. To investigate the time course of these correlations, Spearman’s Rho was calculated at each time point. For condition-level alpha decoding, we investigated correlation between memory and decoding accuracy on 1,000-2,000 ms, the same time window reported in Fellner et al. (2020).

In item-level time-domain decoding, we investigated the correlations between decoding accuracy and absolute memory score of the same condition, on 0-500 ms and 500-3000 ms, respectively. To link item-level decoding with condition level inhibitory control theta power change, we calculated correlation between decoding accuracy difference between Think and No-Think on 420-600 ms, and theta power difference between Think and No-Think on 200-400 ms.

In the High- vs. Low-Suppression Grouping correlation, we calculated correlation between decoding accuracy on 300-680 ms and No-Think minus Baseline Detail memory score, to be consistent with the grouping measure.

#### Highvs. Low-Suppression Grouping

We divided 40 participants into Highvs. Low-Suppression Groups, with 20 participants in each group based on the median split of No-Think-minus-Baseline Detail scores ranking. We used Detail because it captured both variability and suppression effects to a greater extent than did Identification (limited variability since it was a dichotomous measure) and Gist (did not show suppression effect). Pre-TNT learning was not different between Think and No-Think in either group (*p*s > .116).

### Quantification and Statistical Analysis

#### Behavioral Analyses

We conducted separate one-way ANOVAs with three within-subject conditions (Think vs. No-Think vs. Baseline) on the percentage of Identification, Gist, and Details. We then examined the suppression-induced forgetting effect by conducting planned pairwise t-test between No-think and Baseline, with a negative difference (i.e., No-Think minus Baseline) indicating below-baseline, suppression-induced forgetting.

We report findings with *p* < .05 as significant. Within-subject analyses of variance (ANOVAs) are reported with Greenhouse-Geisser corrected *p*-values whenever the assumption of sphericity was violated. We report Cohen’s dz as effect size given our within-subject design (Lakens, 2013).

#### Condition-/Item-level Decoding with Time Domain EEG

Following the statistical analysis procedure (Bae & Luck, 2018), decoding accuracy at each time point (on 0-3000 ms) was compared to chance level by one-tailed paired t-test. Multiple comparisons were controlled by non-parametric cluster-based Monte-Carlo procedure. Specifically, a null distribution was constructed by assigning trial level classification results to random classes (as if the classifier has no knowledge of actual information), and then timepoint-by-timepoint t-tests were performed to obtain a maximum summed t-value of continuous significant time cluster, which then repeated for 1,000 times. The resulting null distribution contained 1,000 summed t-values, which would be the distribution of the cluster summed ts when there is no true difference between decoding results and chance level. Both the cluster *α* and the *α* to obtain critical values from the permutation null distribution were set at 0.05 (on the positive tail, one-tail against chance).

The between-condition comparison of decoding accuracy along time were similar, except that the null distribution was constructed by randomly assigning condition labels to trial level classification results with two-tail repeated measure t-test and clusters were obtained on positive/negative tails, respectively. Thus, the critical values from the permutation null distribution were at 2.5% on the negative clusters null distribution and 97.5% on the positive clusters null distribution.

#### Channel Searchlight Decoding

We compared channel searchlight topographies between item-level decoding in Think and Nothink with a two-tailed paired-sample t test at each channel. The multiple comparisons were controlled by cluster correction of channel neighbour clusters in Fieldtrip (Oostenveld et al., 2011). The neighborhood was defined as in the channel searchlight analysis. Cluster *α* was set at 0.05. Observed clusters were compared to null distribution on positive/negative tails respectively.

#### Condition-/Item-level Decoding with Time-Frequency Domain EEG

The statistical analyses for time-frequency domain decoding were similar to those of time domain decoding, except that here clusters were calculated in a 2-D matrix instead of on a 1-D time axis, and the cluster *α* was set at 0.05. Also, observed clusters were compared to the null distribution clusters of the same rankings. The statistical comparison of a single timefrequency decoding was performed against chance level (one-tailed), and that of the difference between two time-frequency decoding was performed against 0 (two-tailed). Theta (4-8 Hz) and alpha (9-12 Hz) decoding were assessed after averaging across the corresponding frequency bin.

#### Time Frequency Analyses

Early theta power at each electrode was compared between No-Think and Perceptual Baseline after averaging on 200-400 ms across 4 to 8 Hz, and then cluster corrected according to electrode positions in Fieldtrip (Oostenveld et al., 2011). The suppression-associated reduction of theta and alpha power on later time window was examined by averaging on 500-3000 ms across 4-8 Hz (theta) and 9-12 Hz (alpha), and then compared between No-Think and Think/Perceptual Baseline with neighbour cluster correction in Fieldtrip. The channel neighbours were defined in the same way as in channel searchlight analysis.

#### Correlation Analyses

The cluster correction for correlation time course was performed: we first transformed Spearman’s Rho back to t-values to obtain the observed time-course clustered t-values and a null distribution. The null distribution was obtained by randomizing labels of the two variables of interest before calculating the Spearman’s Rho and corresponding t-value. The cluster alpha was set as 0.05, and the observed clusters were calculated for positive and negative clusters respectively. The critical values of null distribution were at the 2.5% on both tails. The comparison between 2 correlation coefficients was conducted through a two-sided z test controlling for dependence (Lenhard & Lenhard, 2014).

#### Highvs. Low-Suppression Groups Comparison

Decoding accuracy at each time point on 0-3000 ms was compared between Highand Low suppression groups using two-tail independent t-test. The null distribution was constructed by randomly assigning group labels to each subject before by-timepoint t-test, to obtain the max summed-t of continuous significant time cluster when group labels are randomized, which repeated for 10,000 times. The resulting 10,000 summed-t values would be the null distribution when no true difference exists between the two groups. Critical values from the permutation null distribution were at 2.5% on the negative clusters null distribution and 97.5% on the positive clusters null distribution (two-tail, *α*s = 0.05).

## Notes

### Competing Interest Statement

The authors have declared no competing interest.

