## Supplemental Figures 1-3 for "Observing the suppression of individual aversive memories from conscious awareness"

**Supplemental Information**

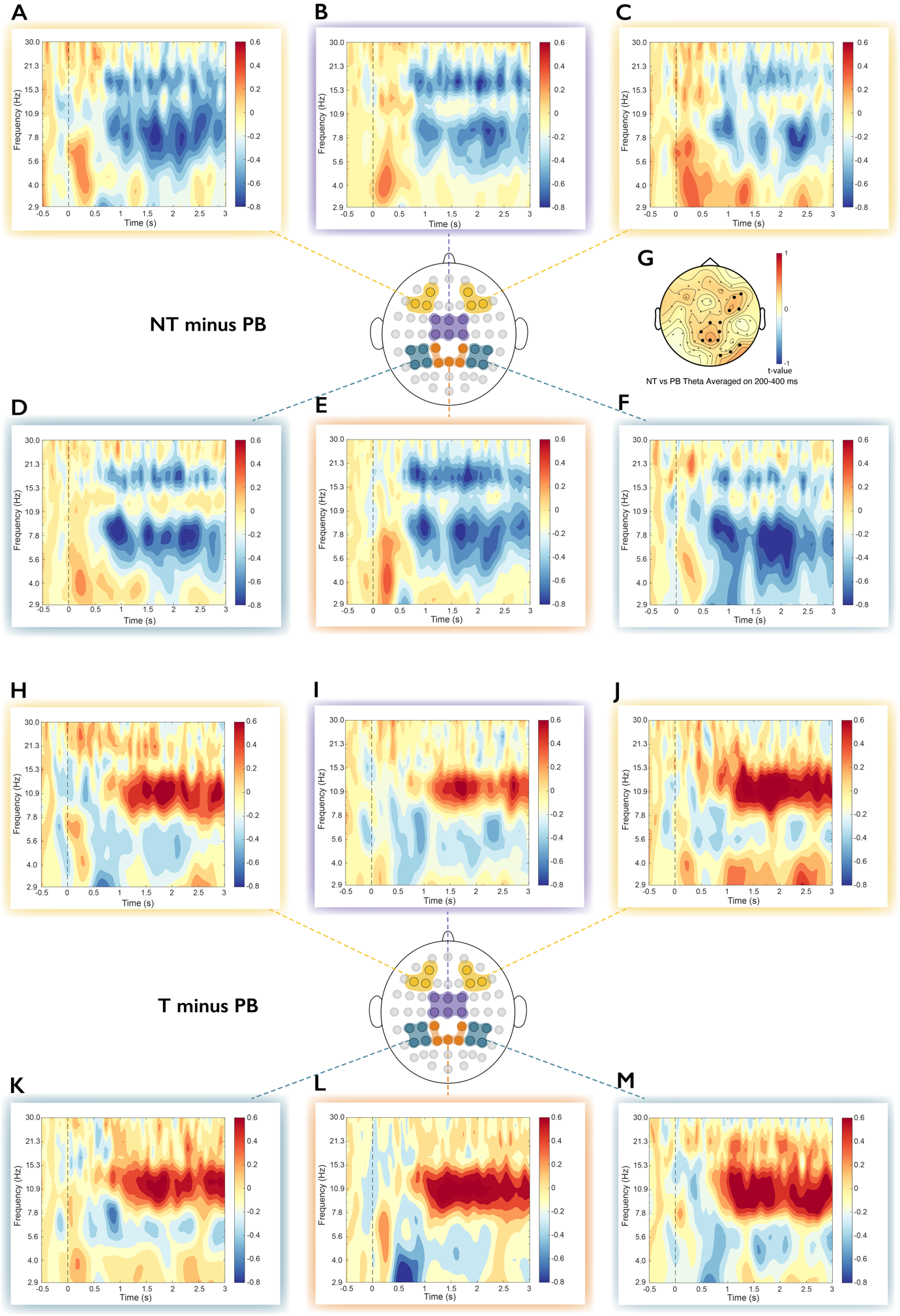

Figure S1. Time Frequency Results.

(A, C) No-Think minus Perceptual Baseline TFRs averaged at channels over left and right Prefrontal regions: left-prefrontal region includes AF3, F3, F5; right- prefrontal region includes AF4, F4, F6.

(B) No-Think minus Perceptual Baseline TFRs averaged at channels over Frontal-central region (Fz, F1, F2, Cz, C1, C2).

(D-F) No-Think minus Perceptual Baseline TFRs averaged at channels over three Parietal regions: left-parietal (CP3, CP5, P3, P5), central-parietal (CP1, CP2, Pz, P1, P2), and right-parietal (CP4, CP6, P4, P6).

(G) Theta oscillations averaged on 200-400 ms increased in No-Think vs. Perceptual Baseline over prefrontal- and central parietal regions. Electrodes with significant difference are highlighted (cluster-corrected).

(H, J) Think minus Perceptual Baseline TFRs averaged at channels over left and right Prefrontal regions: left-prefrontal region includes AF3, F3, F5; right- prefrontal region includes AF4, F4, F6.

(I) Think minus Perceptual Baseline TFRs averaged at channels over Frontal-central region (Fz, F1, F2, Cz, C1, C2).

(L-M) Think minus Perceptual Baseline TFRs averaged at channels over three Parietal regions: left-parietal (CP3, CP5, P3, P5), central-parietal (CP1, CP2, Pz, P1, P2), and right-parietal (CP4, CP6, P4, P6).

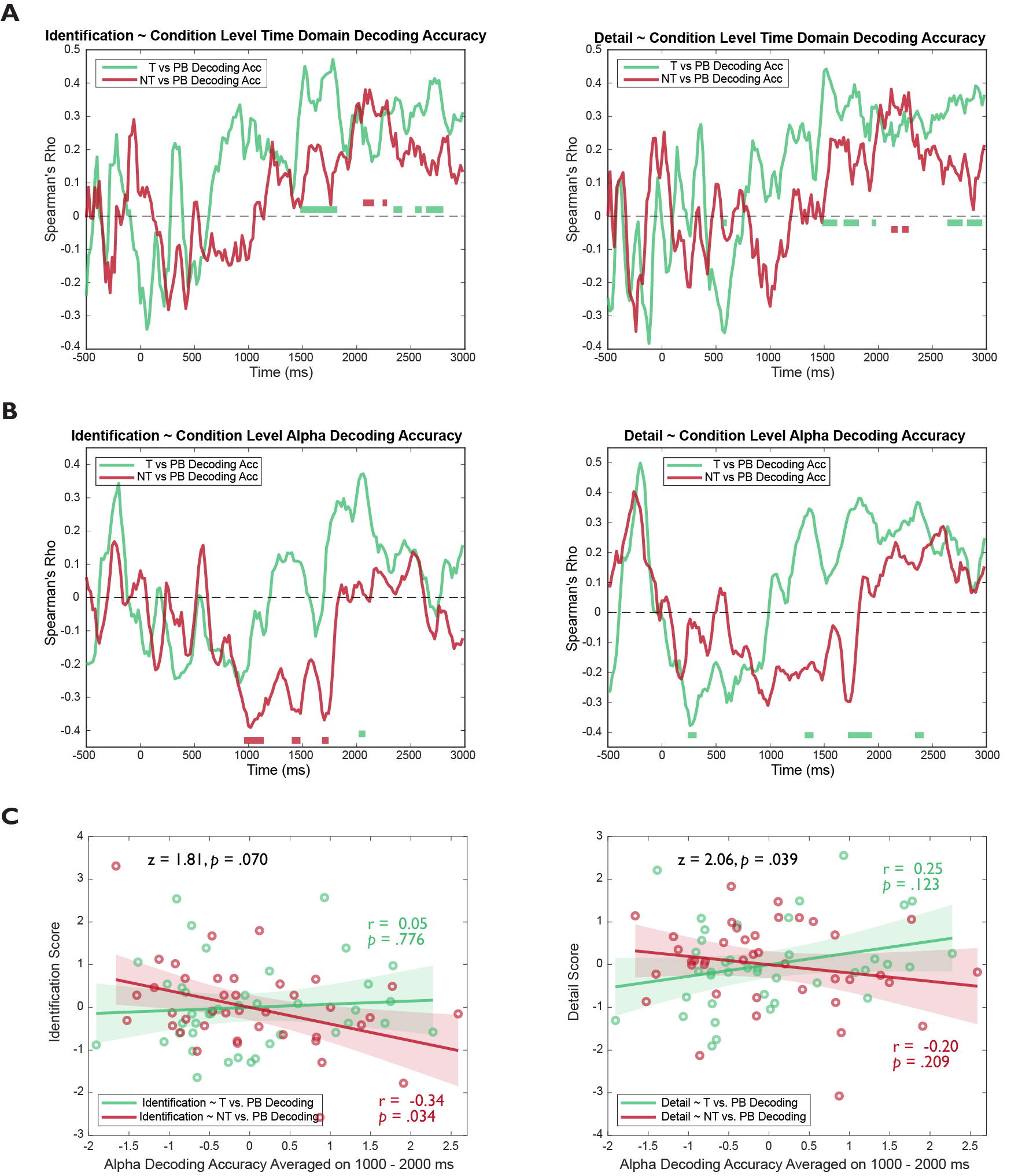

Figure S2. Correlation Between Condition Level Decoding Accuracy and Memory.

(A) Correlation timeline between condition-level time domain decoding accuracy and memory (from left to right: *Identification*, *Detail*). Memory is normalized within each participant, by subtracting participant’s *BL* memory from *T*/*NT*, then divided by *BL*. *Identification* and *Detail* positively correlated with *T* vs. *PB* decoding accuracy on later time windows.

(B) Correlation timeline between condition-level alpha oscillation decoding accuracy and memory (from left to right: *Identification*, *Detail*). Alpha oscillation decoding accuracy was the averaged decoding accuracy from 8.7-12.2 Hz in time-frequency decoding. *NT* vs. *PB* alpha decoding accuracy was found to be negatively correlated with *Identification* and *Gist* on around 1,000-2,000 ms.

(C) Scatter plots of correlations between condition-level alpha oscillation decoding accuracy averaged on 1,000-2,000 ms and memory (from left to right: *Identification*, *Detail*).

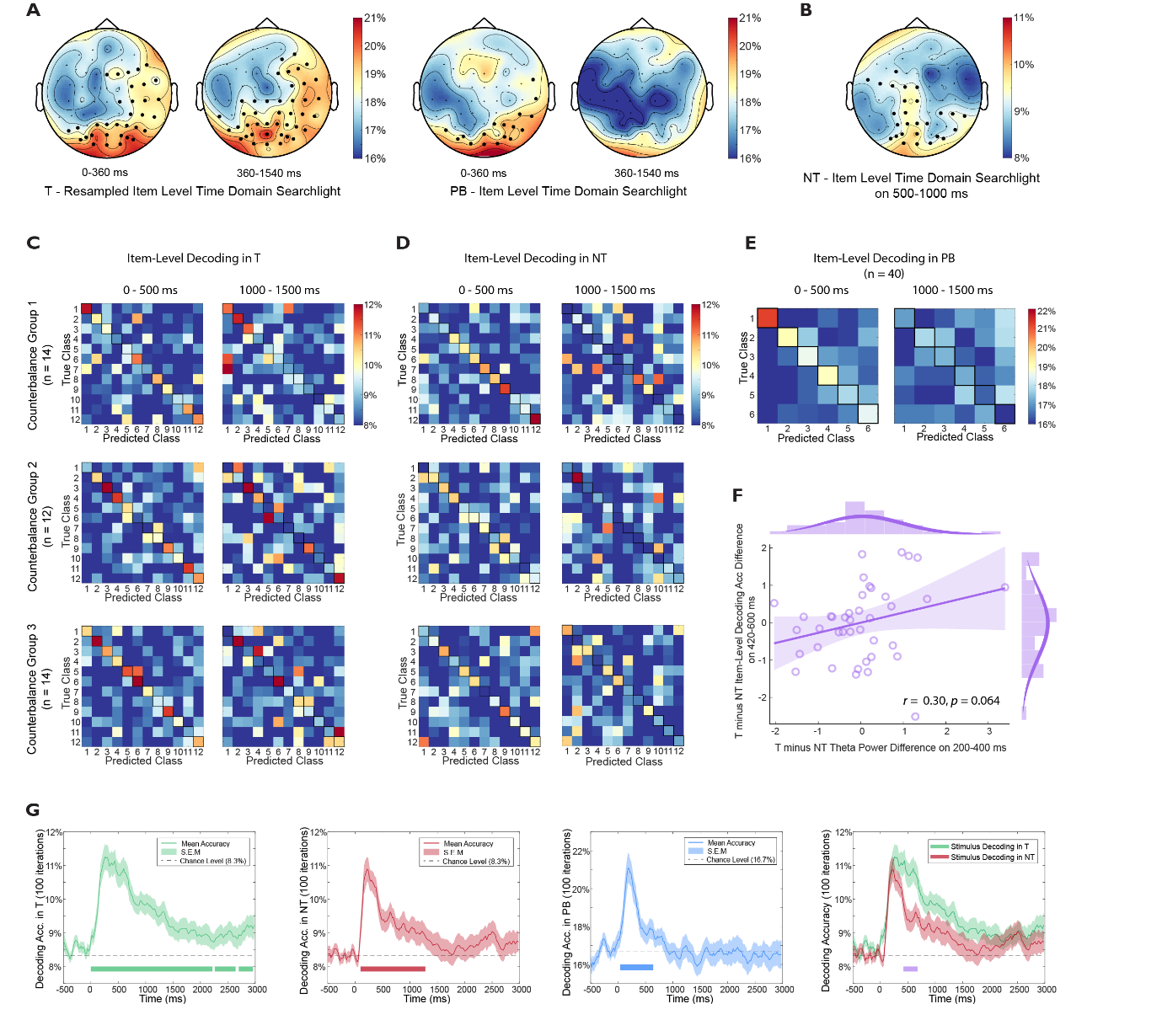

Figure S3. Item-Level Time Domain EEG Decoding Confusion Matrices and Individual Difference Results.

(A) Time-domain item-level searchlight on 0-360 ms showed an occipital distribution in Think (resampled) and Perceptual Baseline; on 360-1540 ms, Think showed a right-prefrontal-parietal-occipital distribution, while no electrodes could decode items in Perceptual Baseline.

(B) Time-domain item-level searchlight in No-Think showed an occipital-parietal distribution on 500-1000 ms.

(C-E) Confusion matrices of item-level decoding accuracies averaged on 0-500 ms and 1000-1500 ms. In *T* & *NT* (C, D), items were divided into three counterbalance groups and their confusion matrices were plotted separately. No counterbalance was performed for Perceptual Baseline (E).

(F) Significant reduction of item-level decoding accuracy in No-Think (vs. Think, on 420-600 ms) positively correlated with enhanced theta power in No-Think (vs. Think) on 200-400 ms.

(G) Item-level time domain decoding was validated with 100 iterations. The results were highly similar to those reported in main text (using 10 iterations). Moreover, the comparison between *T* and *NT* item level decoding found a significant cluster on 420-680 ms, which was also consistent with 10-iteration results.

Electrodes with significant decoding accuracies are highlighted (cluster-corrected).
